# Increased light availability enhances tolerance against ocean acidification stress in *Halimeda opuntia*

**DOI:** 10.1101/2020.10.09.333799

**Authors:** Zhangliang Wei, Chao Long, Yating Zhang, Yuanzi Huo, Fangfang Yang, Lijuan Long

## Abstract

Although the adverse impacts of ocean acidification (OA) on marine calcifiers have been investigated substantially, the anti-stress abilities regulated by increased light availability are unclear. Herein, the interactive effects of three light levels combined with two *p*CO_2_ concentrations on the physiological acclimation of the calcifying macroalga *Halimeda opuntia* were investigated using a *p*CO_2_–light coupling experiment. The results indicate that OA exhibits an adverse role in influencing algal growth, calcification, photosynthesis and other physiological performances in *H. opuntia.* The relative growth rate in elevated *p*CO_2_ significantly declined by 13.14%–41.29%, while net calcification rates decreased by nearly three-fold under OA. Notably, increased light availability could enhance stress resistance by the accumulation of soluble organic molecules, especially soluble carbohydrate, soluble protein and free amino acids, and in combination with metabolic enzyme-driven activities alleviated OA stress. Carotenoid content in low light conditions accumulated remarkably and rapid light curves for relative electron transport rate was significantly enhanced by increasing light intensities, indicating that this new organization of the photosynthetic machinery in *H. opuntia* accommodated light variations and elevated *p*CO_2_ conditions. Taken together, the results describe stress resistance by the enhancement of metabolic performance in marine calcifiers to mitigate OA stress.

**One sentence summary:** Increased light availability enhances stress resistance in *Halimeda opuntia* by the accumulation of soluble organic molecules and enzyme-driven activities to alleviate ocean acidification stress.

**Credit authorship contribution statement:** Fangfang Yang and Lijuan Long conceived and designed the experiments. Zhangliang Wei performed the experiments and wrote the paper. Yuanzi Huo analyzed the data, while Chao Long and Yating Zhang contributed materials and analysis tools. Lijuan Long agrees to serve as the author responsible for contact and communication.

**Highlights:**

1. Elevated *pCO_2_* adversely affects the physiological performance of *Halimeda.*
2. Moderately high light increases soluble organic molecules and enzymatic-driven activities.
3. Increased light availability enables *H. opuntia* to alleviate the negative effects of ocean acidification.

## 1. Introduction

The atmospheric concentration of carbon dioxide (CO_2_) has been rising rapidly over the past 250 years due to large-scale use of fossil fuels and deforestation, and is predicted to exceed 1000 ppmv by the end of 2100 unless anthropogenic CO_2_ emissions are effectively managed (Fabry et al., 2008; IPCC, 2013). Concurrently, the oceans have already absorbed approximately 25 million tonnes of emitted CO_2_ every day. This has stimulated a decrease in pH (~0.1 unit) in the surface layer of oceans and is projected to reach a mean of pH 7.8 by the end of 2100 (IPCC, 2013). CO_2_-induced ocean acidification (OA) has received substantial attention and believed to cause combined interactive effects on marine ecosystems (Orr et al., 2005; Doney et al., 2009; Cornwall et al., 2013). In terms of the impact of OA on marine calcifiers, including calcifying macroalgae, the main focus has been the possible direct impacts of decreasing the availability of carbonate ions (CO_3_^2−^) and carbonate saturation (Moya et al., 2012; Hofmann et al., 2014; Wei et al., 2020a). In common with previous studies, OA generally has a negative effect on the biomineralization process caused by reduced seawater pH and aragonite saturation state (Ω) (Ries et al., 2009; 2010; Comeau et al., 2017). Campbell et al. (2016) have reported that under elevated *p*CO_2_ conditions (2400 ppmv), calcification rates of two calcifying macroalgae species *(Halimeda opuntia* and *H*. *simulans*) declined by 32% and 49%, respectively. These conclusions have signified that the consequences of OA are expected to reduce the distribution of calcifiers, CaCO_3_ stocks and biomass, which will result in various changes in habitat structure and ecological function for marine ecosystems worldwide (Gao et al., 1993; Kleypas and Yates, 2009). However, variable effects have been reported because some species display relatively minor responses to OA. Peach et al. (2017) noted that calcification and growth rates varied among six *Halimeda* species, but with no obvious effects at elevated *p*CO_2_ (1300 ppmv). Although species-level variation in response to OA has been documented in rates of calcification and growth in field and laboratory experiments, significant gaps still exist in our knowledge of the metabolic response to OA by calcifiers due to the interactive effects of other dynamic environmental parameters and different meta-analyses (Fabry et al., 2008; Doney et al., 2009; Wei et al., 2020a).

Calcifying macroalgae are considered to be an important part of coral reef ecosystems as their calcium carbonate endoskeletons are laid down and provide vast, three-dimensional structures and shelters for many coral inhabitants (Wei et al., 2020b). In the open ocean, light is another vital environmental factor for the growth and distribution of macroalgae (Zou and Gao, 2010). Variations in light availability can affect algal photosynthesis, carbohydrate synthesis and substance accumulation through photo energy absorbed by light-harvesting complexes (Porzio et al., 2011; Celis-Plá et al., 2017). Previous studies have found that moderate increases in light intensity have a positive influence on the physiological performance of fleshy and calcifying macroalgae (Teichberg et al., 2013; Bao et al., 2019). Teichberg et al. (2013) pointed out that in Curacao, Netherlands Antilles, the growth, photosynthetic performance and tissue composition (total organic carbon and nitrogen) of *H. opuntia* at depths of 5 m were significantly higher than those at 15 m mainly due to differences in the incidence of underwater light. High sunlight in relatively shallow marine environments provides sufficient solar energy for calcifying macroalgae to uptake bicarbonate (HCO_3_^−^) through carbon concentrating mechanisms (CCMs) (Borowitzka and Larkum, 1976; Koch et al., 2013). As CO_2_ dissolved in seawater can increase the concentration of HCO_3_^−^, which is used for the assimilation of biogenic calcium carbonate (Orr et al., 2005), it is essential to conduct experiments that combine OA scenarios with other factors such as light availability (Hofmann et al., 2014).

The genus *Halimeda* (Bryopsidales) is composed of calcified green segments that play a major role as carbonate sediment producers and is distributed widely in tropical and subtropical marine ecosystems (Payri, 1988; Sinutok et al., 2012; Wizemann et al., 2015). It has been estimated that *Halimeda* species produce 0.15–0.40 Gt of CaCO_3_ m^−2^ year^−1^, accounting for around 8.0% of the total calcium production of coral reef ecosystems (Milliman, 1993; Hillis, 1997). Thus, *Halimeda* species are considered a carbon sink for the long-term carbon storage of atmospheric CO_2_ emissions (Kinsey and Hopley, 1991; Rees et al., 2007). The calcification of this genus is a light-driven process, requiring functional chloroplasts in superficial utricles to remove CO_2_ from the seawater into interfilamental spaces (Borowitzka and Larkum, 1976). Substantial studies have shown that OA brings negative influences on growth, calcification, photosynthesis and other aspects of the physiological performance of *Halimeda* (Hofmann et al., 2014; Wei et al., 2020a). Under elevated *p*CO_2_ stress, malondialdehyde (MDA) content in these calcifying macroalgae significantly increases. MDA is a secondary end product of oxidative lipid degradation in cells and has deleterious effects on protein synthesis and nucleic acid reactions (Xiong et al., 2002; Hemm et al., 2004; Wei et al., 2019; 2020a).

It is essential that marine macroalgae make suitable modifications to deal with OA stress. Functional soluble organic osmolytes, such as proline, soluble carbohydrate (SC), soluble protein (SP), and free amino acids (FAA), are secreted in abundance and protect the integrity of cellular structures, as well as other defense-related enzymes systems (Chen et al., 2017; 2018; Wei et al., 2020a). Hofmann et al. (2014) suggested that under OA conditions (pH 7.75±0.05), the external carbonic anhydrase activity (eCAA) of *H. opuntia* significantly increased to enhance photosynthetic performance and carbon accumulation abilities. However, it is still not known whether marine calcifying organisms will be able to tolerate future conditions, as the cumulative effects of continued OA drive these calcifying species closer to the limits of their physiological tolerances (O’Donnell et al., 2010; Koch et al., 2013). A moderate increase in light availability could enhance the metabolic activities of *Halimeda* species in response to OA stress, indicating its important role in mitigating adverse influences on these carbonate producers (Teichberg et al., 2013; Wei et al., 2020).

The South China Sea is a tropical sea and is one of the most multifarious coral community habitats (Morton and Blackmore, 2001). The aragonite-depositing species *Halimeda opuntia* can be found widely in this sea area (Wei et al., 2020b). Taking specific growth locations and depth ranges into consideration, *H. opuntia* experiences varied sunlight irradiance conditions during the daytime (El-Manawy and Shafik, 2008; Peach et al., 2017). Although our previous study found that increased light intensities have positive effects on the physiological performance of *Halimeda*, it will be necessary to conduct further study to explore the key anti-stress mechanisms of this calcifying species in response to OA under light regulation. A detailed description of anti-OA stress mechanisms will provide an important scientific basis for the prediction of spatial distribution in different seawater depths and population protection under global climate changes.

## 2. Materials and methods

### 2.1 Sample collection

Samples of *H. opuntia* were collected by SCUBA in a back reef lagoon in the South China Sea (17.03°–17.07° N, 111.28°–111.32° E) at depths ranging from 12–15 m in July 2019. All individuals were carefully detached from the sandy substrate. After collection, algal thalli were immediately incubated in aquaria with a flowing system receiving ambient seawater in the ship and transported back to the marine research laboratory in Sanya, China.

For acclimation, the algae were reared in an aerated mesocosm tank (1,200 L) with a constant supply of fresh sandy-filtered seawater (~30 L min^−1^) in the laboratory for two weeks. Ambient seawater conditions consisted of 27.0°C, 32 ppt, pH 8.1 and 80 μmol photons m^−2^ s^−1^ with a 12:12 h day/night cycle. After two weeks acclimation, thalli of *H. opuntia* turned a healthy green and subsequent experiments could be undertaken (Hofmann et al., 2014).

### 2.2 Experimental design and treatments

For the laboratory trial, healthy thalli of *H. opuntia* were randomly assigned to 2 × 3 factorial coupling treatments with three light intensities and two seawater *p*CO_2_ levels to determine the mixed effects on the physiological performance of *H. opuntia* for four weeks (10^th^ November to 8^th^ December 2019). Individuals with a fresh weight (FW) of 140–150 g were assigned to each aquarium (30 L) and six treatments consisted of 18 aquaria due to three biological replications (Table 1). Filtered seawater was changed twice a week. OA (elevated *p*CO_2_) was achieved by bubbling different concentrations of mixed CO_2_ and ambient air (CE100C, Wuhan Ruihua Instrument and Equipment Ltd., Wuhan, China). High *p*CO_2_ (HC) was set at 1400 ppmv, which is the predicted level under extreme conditions at the end of this century (Caldeira and Wickett, 2005). The other *p*CO_2_ level of 400 ppmv (LC) was considered to be the current average seawater *p*CO_2_ concentration. Light intensities were set for three levels of 30 (low, LL), 150 (middle, ML) and 240 (high, HL) μmol photons m^−2^ s^−1^, which were created using full-spectrum fluorescent aquarium lamps (Giesemann, Nettetal, Germany) and monitored by an optical quantum probe (ELDONET Terrestrial Spectro-radiometer, Frankfurt, Germany). Although the light intensities selected in the present study were lower than the sampling site (300–500 μmol photons m^−2^ s^−1^), previous research has revealed that the saturation light (I_k_) of *Halimeda* species in situ typically ranged from 30 to 250 μmol photons m^−2^ s^−1^ (Beach et al., 2003; Hofmann et al., 2014; Campbell et al., 2016).

**Table 1.** Factorial design used to test the effects of elevated *p*CO_2_ at three light levels.

| Treatments<br>($p\text{CO}_2 \times \text{Light}$) | Seawater $p\text{CO}_2$<br>(ppmv) | Light intensities<br>($\mu\text{mol photons} \cdot \text{m}^{-2} \text{s}^{-1}$) |
| --- | --- | --- |
| HC-LL | 1400 | 30 |
| HC-ML | 1400 | 150 |
| HC-HL | 1400 | 240 |
| LC-LL | 400 | 30 |
| LC-ML | 400 | 150 |
| LC-HL | 400 | 240 |

### 2.3 Monitoring experimental conditions

To ensure the consistency of seawater between treatments, seawater temperature and salinity were monitored daily by a calibrated handle YSI meter (YSI Professional Plus, Yellow Springs, OH, USA). The pH (NBS scale) was determined using an S220 pH electrode (Mettler Toledo, Switzerland). Before the measurement, the pH electrode was calibrated by a standard buffer solution (pH 4.003, 6.864 and 9.182). Total alkalinity (TA) was measured using the Gran titration method (Metrohm 877 Titrino Plus) and the specific protocol was described by Dickson et al. (2007). All measurements were conducted under 25.0°C and completed within 24 h of sampling. Dissolved inorganic carbon components (CO_2_, CO_3_^2−^ and HCO_3_^−^) and aragonite saturation state (Ω_Arag_) were calculated by salinity, temperature, pH_NBS_ and TA with the Excel program CO2SYS (Pierrot et al., 2006).

### 2.4 Measurement of growth and calcification rates

The FW (g) of *H. opuntia* was measured at the beginning and the end of the experiment. The growth rate was calculated using the following formula:

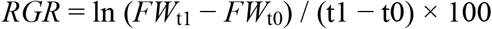

where *RGR* is the relative growth rate (*RGR*) (% d^−1^); *FW*_t1_ and *FW*_t0_ are the fresh weights at time t1 and t0 days, respectively.

At the end of the experiment, the net calcification rate (*G*_net_) was calculated by the TA anomaly technique according to the description by Hofmann et al. (2014), which is based on the equation that every mole of formed CaCO_3_ reduces the TA by two equivalents.

### 2.5 Chlorophyll fluorescence and pigment contents

The photosynthetic performance of *H. opuntia*, including maximum quantum yield of photosystem II (*F*_v_/*F*_m_) and relative electron transport rate (rETR), were measured at the end of the experiment via pulse amplitude modulated (PAM) fluorometry with a DIVING-PAM II fluorometer (Heinz Walz GmbH, Effeltrich, Germany). On the sampling date, haphazard mature *Halimeda* thalli in each treatment were selected and adapted in darkness for nearly 20 min before measurement. The rapid photosynthesis-light curves were determined under different photosynthetically active photon flux levels by 1 min intervals. The maximum relative electron transport rate (rETRmax) was calculated from a nonlinear regression analysis using the model described by Eilers and Peeters (1988). Approximately 100.0 mg FW segment was sampled for photosynthetic pigments. Chlorophyll a (Chl-*a*) and carotenoids (Car.) were calculated according to the method reported by Wellburn (1994).

### 2.6 Determination of organic carbon and nitrogen content

Total tissue organic carbon (TCorg) and total nitrogen (TN) contents were measured after four-week experimental periods. Samples were cleaned of epiphytes, detritus and sediment by washing with filtered seawater 3–5 times. To determine TCorg, dry tissue samples were acidified by 1 N HCl until no air bubbles were observed. Subsequently, all samples were dried at 60°C until no change in dry mass weight was detectable and were finely ground to a powder with an automatic grinding mill (FSTPRP-24, Jingxin, Shanghai, China). TC_org_ and TN were measured using a CHN Elemental Analyzer (Flash EA300, Thermo Scientific, Milan, Italy).

### 2.7 Measurement of enzyme activity

At the end of the experiment, 200.0 mg fragments were sampled to measure the eCAA and nitrate reductase activity (NRA), respectively. The eCAA of *H. opuntia* was measured using the electrometric technique described by Haglund et al. (1992), while NRA was determined by the method reported by Corzo and Niell (1991).

### 2.8 Soluble carbohydrate, protein and free amino acid content

Approximately 200.0 mg FW fragments were finely ground with a mortar and pestle with 10 ml distilled water. Afterwards, the extract was centrifuged at 5, 000 rpm for 10 min (Eppendorf centrifuge 5810R, Hamburg, Germany) and then the supernate was used to determine SC and SP using an ultraviolet spectrophotometer (UV 530, Beckman Coulter, Brea, USA). The SC content was determined using the phenol-sulfuric acid method (Kochert, 1978a). The SP content was measured by Coomassie Brilliant Blue G-250 dye combination and standardized with bovine albumin (Kochert, 1978b).

To determine FAA, 200.0 mg FW algae were sampled and finely ground in 4.5 ml of PBS buffer (pH 7.4). The homogenate was centrifuged at 5, 000 rpm for 10 min. The supernatant was collected and preserved at 4.0°C. The FAA content was determined using an algal FAAs ELISA Kit (Mlbio, China) following the manufacturer’s instructions.

### 2.9 Statistical analysis

All results in this study are presented as a triplicate mean ± standard deviation (SD). Origin 8.0 software was used to create the figures and data analyses were processed using Minitab 16.0 software. All data used in the univariate analyses were assessed for homoscedasticity and normality. A multivariate analysis of variance (MANOVA) was conducted to analyze the effects of *p*CO_2_ and light as independent factors. Tukey’s honest significance difference (HSD) was used to identify significant differences between treatments and *P* values were significant at the 95% confidence level.

## 3. Results

### 3.1 Seawater conditions

During the experimental period (displayed in Table 2), the average pH within high (1400 ppmv, HC) and control (400 ppmv, LC) CO_2_ treatments was 7.72 and 8.15, respectively. Mean TA in LC (2350 μmol kg^−1^) and HC (2336 μmol kg^−1^) seawater did not differ. However, elevated *p*CO_2_ significantly altered the carbon chemistry parameters in seawater (*P* < 0.01). Compared with LC, the average concentrations of CO_2_ and HCO_3_^−^ in HC increased from 11.66–12.81 to 39.10–39.81 μmol kg ^−1^ and 1782.9–1828.8 to 2138.0–2165.3 μmol kg^−1^, respectively, while CO_3_^2^ concentration decreased from 197.2–206.0 to 86.8–95.9 μmol kg^−1^. The daily calculated saturation state of aragonite (Ω_Arag_) in HC ranged from 3.21 to 3.36 compared to 1.42 to 1.59 in LC (Table 2).

**Table 2.** Measured seawater parameters and calculated carbon chemistry via CO2SYS in each treatment (mean ± SD, n = 3)

| Treatments | pH | TA<br>($\mu\text{mol kg}^{-1}$) | CO<sub>2</sub><br>($\mu\text{mol kg}^{-1}$) | HCO<sub>3</sub><sup>-</sup><br>($\mu\text{mol kg}^{-1}$) | CO<sub>3</sub><sup>2-</sup><br>($\mu\text{mol kg}^{-1}$) | $\Omega_{\text{Arag}}$ |
| --- | --- | --- | --- | --- | --- | --- |
| HC-LL | 7.71±0.11 | 2396±19 | 39.10±0.5 | 2165.3±15.4 | 95.9±1.3 | 1.59±0.13 |
| HC-ML | 7.72±0.06 | 2317±22 | 39.57±0.5 | 2138.0±10.5 | 86.8±0.9 | 1.42±0.09 |
| HC-HL | 7.72±0.07 | 2339±14 | 39.81±0.9 | 2156.8±14.8 | 88.1±1.4 | 1.44±0.11 |
| LC-LL | 8.15±0.14 | 2369±12 | 12.23±0.3 | 1788.1±15.2 | 197.6±1.9 | 3.22±0.25 |
| LC-ML | 8.15±0.09 | 2327±15 | 12.81±0.2 | 1828.8±12.7 | 197.2±2.1 | 3.21±0.21 |
| LC-HL | 8.16±0.05 | 2314±21 | 11.66±0.6 | 1782.9±13.2 | 206.0±2.6 | 3.36±0.22 |

### 3.2 Growth and calcification rates

After four weeks of exposure, elevated *p*CO_2_ and light intensities had clear effects on *RGR* and *G*_net_ in *H. opuntia* throughout the experiment (Fig. 1; Table 3). Highest *RGR* and *G*net values were obtained in LC-HL. The *RGR* of *H. opuntia* in elevated *p*CO_2_ significantly declined by 13.14%–41.29%. However, increased light intensity stimulated growth rates at the two CO_2_ concentration levels. In contrast to low light, the *RGR* increased by 19.28%–33.32% and 26.64%–50.42% in middle and high light, respectively (Fig.1). Similarly, the *G*_net_ decreased by nearly three-fold in HC-LL (0.218 ± 0.035 μmol CaCO_3_ g FW^−1^ h^−1^) compared with that in LC-LL (0.625 ± 0.042 μmol CaCO_3_ g FW^−1^ h^−1^). When the light intensity increased, *G*_net_ increased by 50.27% in ML and 26.16% in HL in the control compared to elevated *p*CO_2_ (Fig.1).

**Fig. 1.**
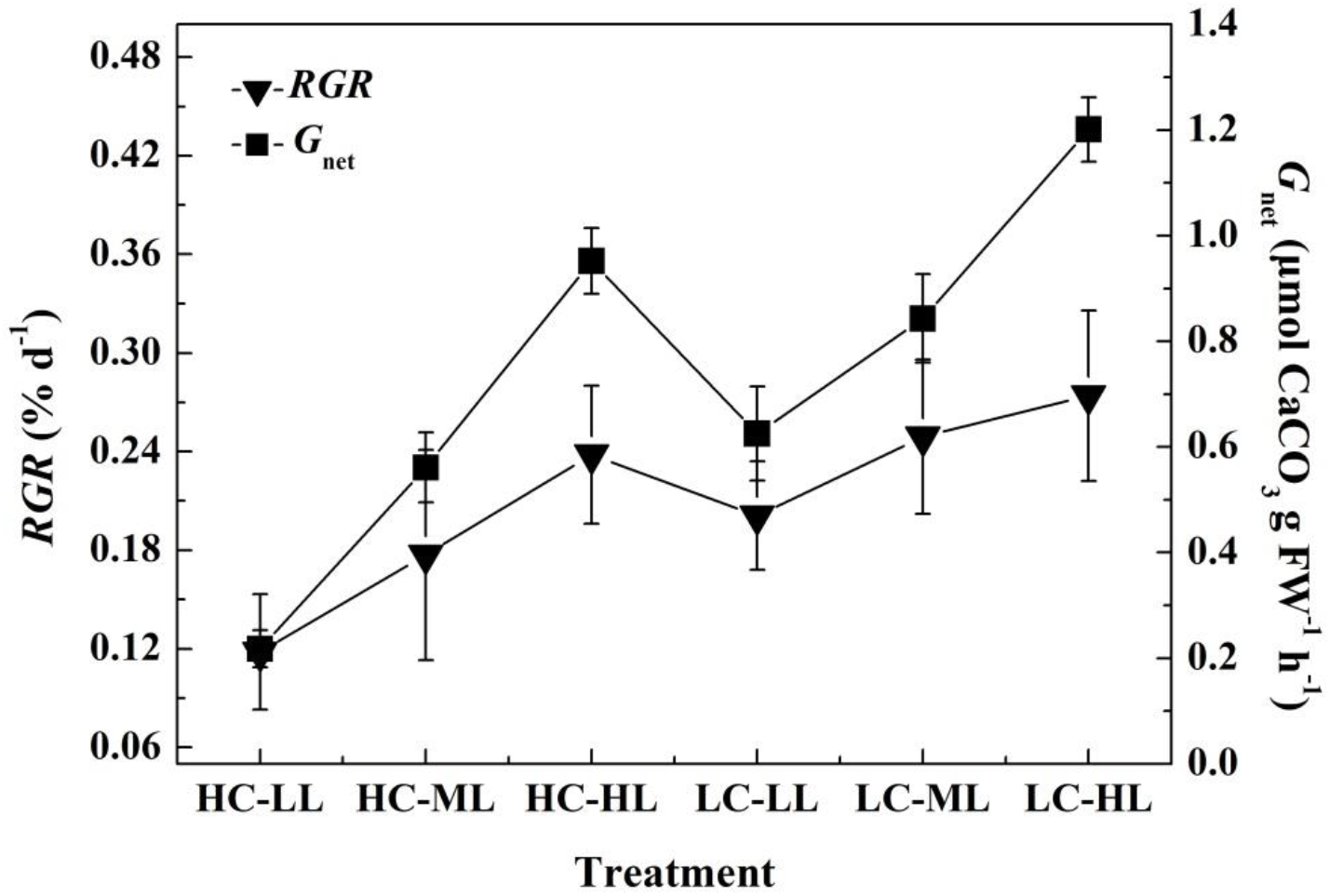
Relative growth rates (*RGR,* % d ^−1^) and net calcification rates (*G*_net_, μmol CaCO_3_ g FW^−1^ h^−1^) in *Halimeda opuntia* (mean ± SD, n = 3) after six *p*CO_2_–light treatments.

**Table 3.**
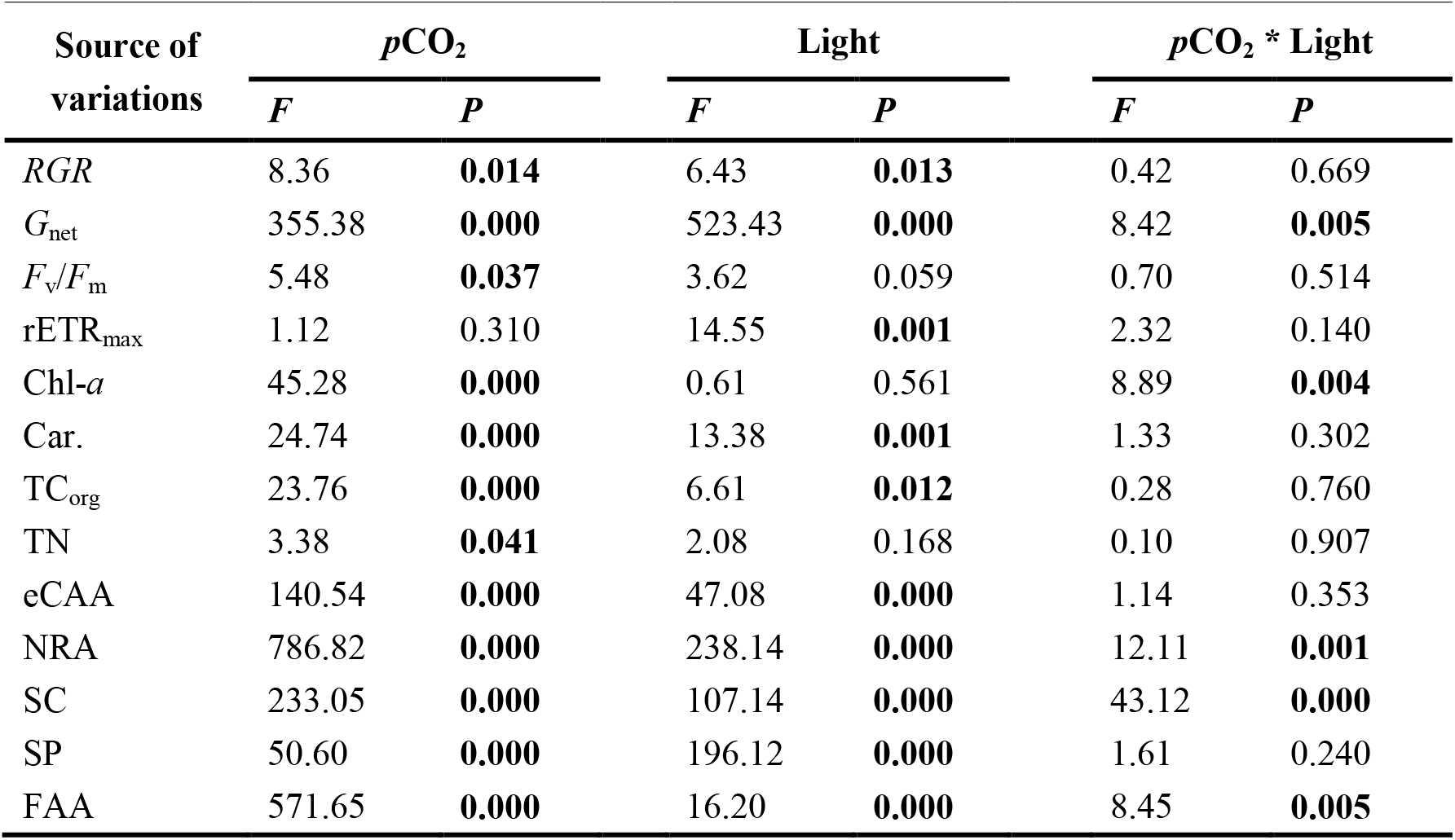
Results of MANOVA tests, including F-ratios and P (95% confidence level), to analyze the effects of two *p*CO_2_ levels crossed with three light intensities on the physiological performance of *Halimeda opuntia*.

### 3.3 Photosynthetic performance and pigment contents

During the experimental period, there were significant effects on rapid light curves with *H. opuntia* rETR in HL conditions, but not significantly affected by elevated *p*CO_2_ in ML nor LL (Fig. 2). Maximum photochemical quantum yield (*F*_v_/*F*_m_) ranged from 0.665 to 0.704 and 0.696 to 0.711 in elevated *p*CO_2_ and control treatments, regardless of the three light levels (Fig. 4a; Table 3). Conversely, the rETR_max_ was only significantly affected by light availability (*P* < 0.01) (Table 3). The rETR_max_ was higher in HL (6.262–6.584 μmol e^−1^ m^−2^ s^−1^) than in ML (5.357–5.694 μmol e^−1^ m^−2^ s^−1^) and LL (5.157–5.674 μmol e^−1^ m^−2^ s^−1^) (Fig. 4b). Notably, *H. opuntia* displayed significant correlation between photosynthetic maximum quantum yield (*F*_v_/*F*_m_) and calcification rate (*G*_net_) (R^2^ = 0.89; *P* = 0.0045) (Fig. 3).

**Fig. 2.**
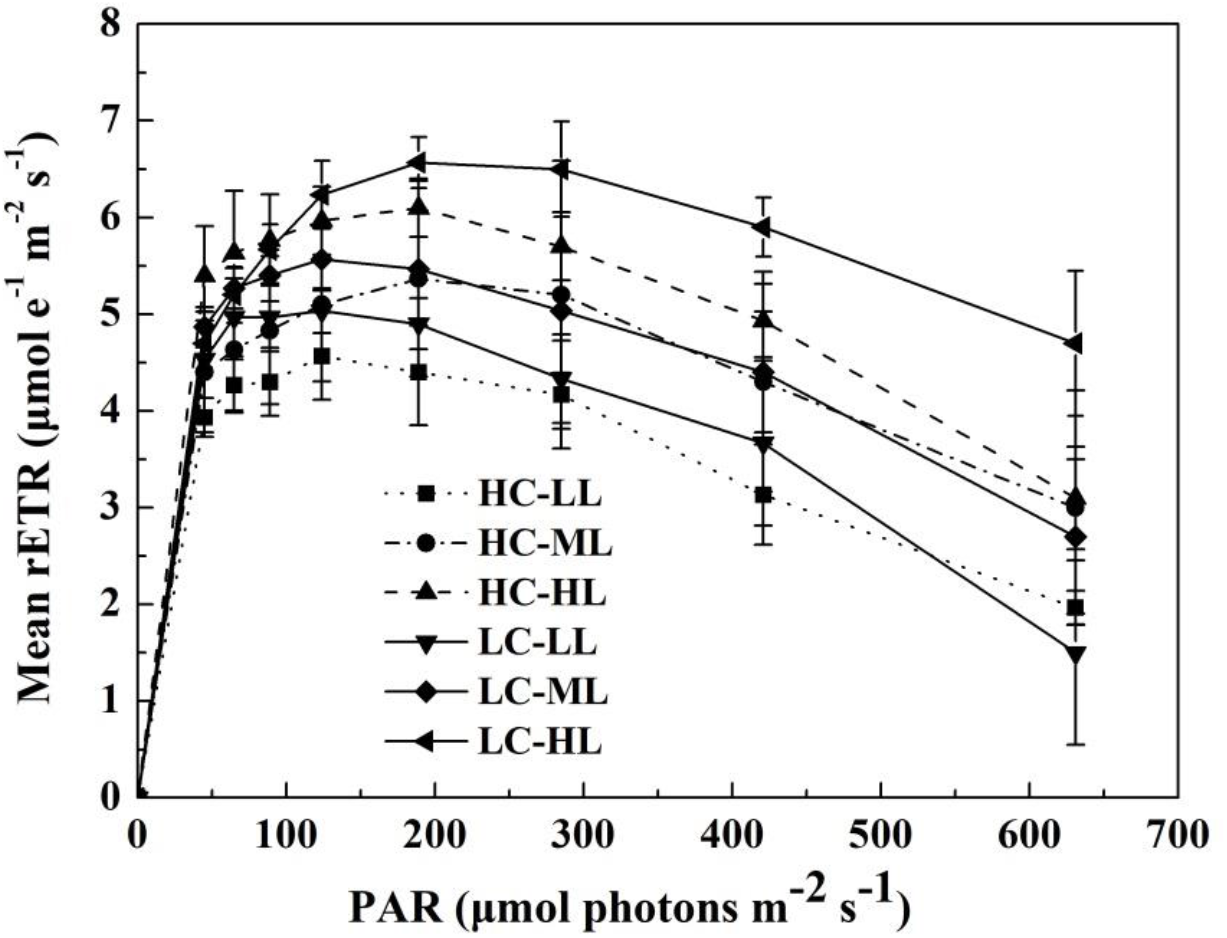
The rapid light curves for electron transport rate (rETR) of photosynthetic active radiation (PAR) of *Halimeda opuntia* incubated in six *p*CO_2_–light treatments.

**Fig. 3.**
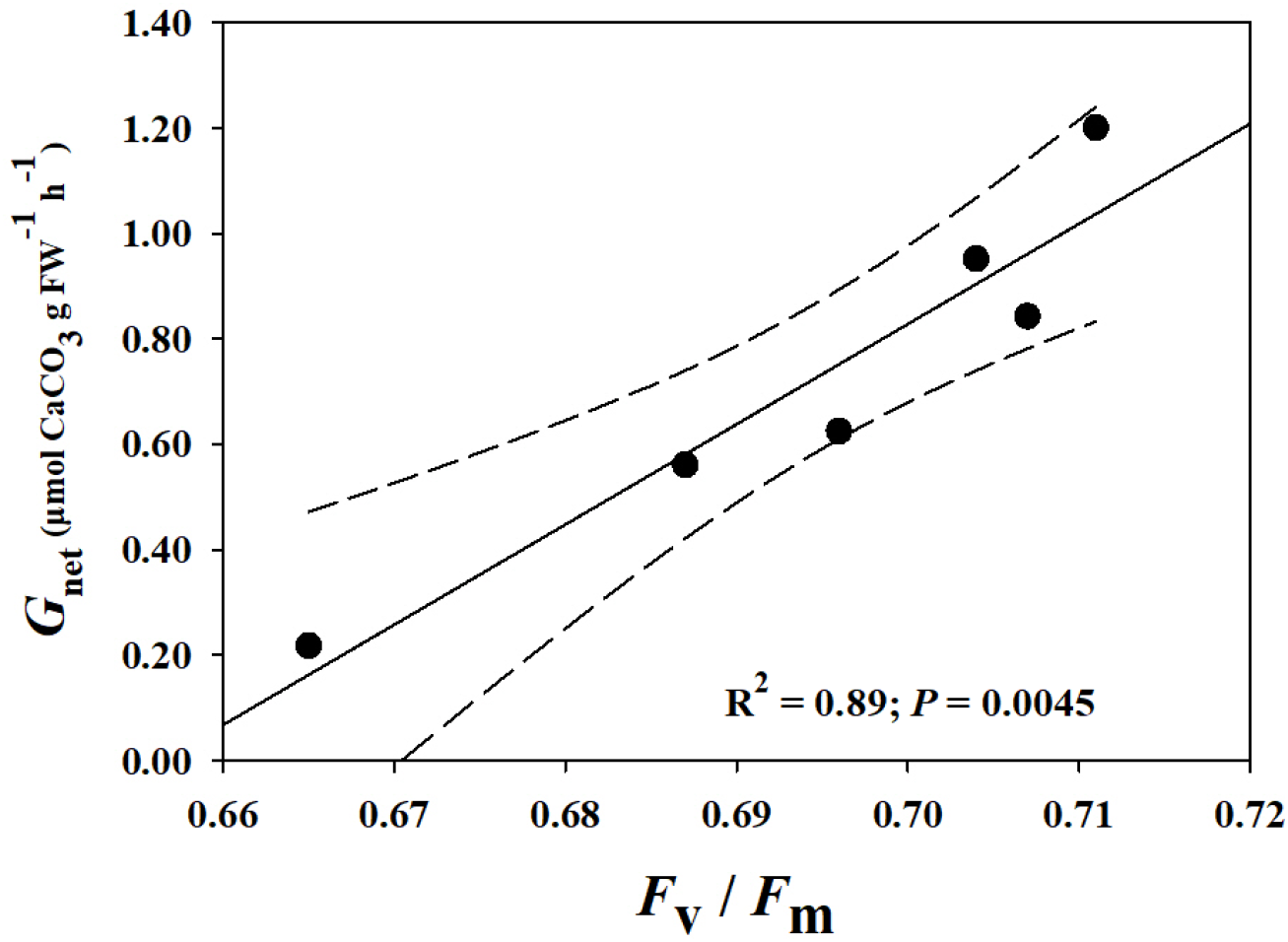
Correlation between photosynthetic maximum quantum yield (*F*_v_/*F*_m_) and calcification rate (*G*_net_, μmol CaCO_3_ g FW^−1^ h^−1^) of *Halimeda opuntia.*

**Fig. 4.**
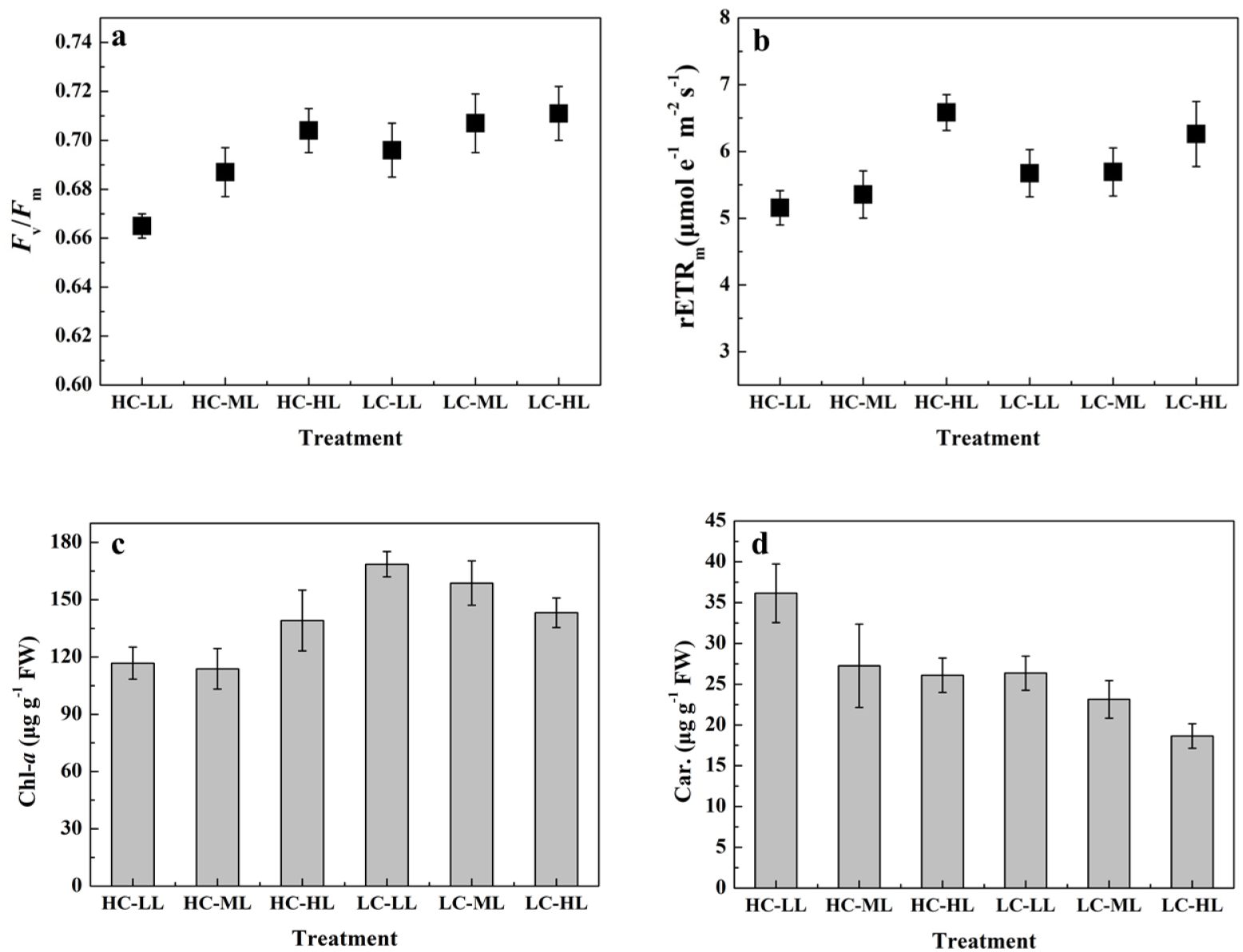
Variations in photosynthetic maximum quantum yield (*F*_v_/*F*_m_) (a), the maximum relative electron transport rate (rETR_m_) (b), pigment contents of chlorophyll (Chl) *a* (μg g ^−1^ FW) (c) and carotenoid (Car., μg g ^−1^ FW) (d) of *Halimeda opuntia* (mean ± SD, n = 3) in six *p*CO_2_-light treatments.

As shown in Table 3, there were significant effects of elevated *p*CO_2_ and light intensities and their interactions on the pigment content of *H. opuntia*. In elevated *p*CO_2_ conditions, the highest Chl *a* content was obtained in HL treatments (139.1 ± 15.9 μg g^−1^ FW), whereas in ambient *p*CO_2_ conditions, the highest Chl *a* content was measured in low light (168.6 ± 6.6 μg g^−1^ FW) (Fig. 4c). Carotenoid content in HC-LL (36.15 ± 3.6 μg g^−1^ FW) was significantly higher than that in LC-HL (18.65 ± 1.5 μg g^−1^ FW), which was affected by elevated *p*CO_2_ and light intensities (*P* < 0.01) (Fig. 4d).

### 3.4 Tissue TC_org_ and TN content

Both elevated *p*CO_2_ (CO_2_ enrichment) and increasing light availability resulted in tissue TCorg accumulation for *H. opuntia* over the four-week experiment (Fig. 5; Table 3). In HC, TC_org_ content significantly increased by 13.78%–17.91% compared with LC (Fig. 5). The highest TC_org_ content (17.64 ± 1.02% DW, dry weight) occurred in HC-HL and the lowest content (13.28 ± 1.34% DW) was measured in LC-LL. TN content exhibited a similar pattern of variation, as the elevated *p*CO_2_ significantly stimulated nitrogen accumulation (*P* = 0.041) (Table 3). However, light variation had no effect on the TN content in *H. opuntia* as shown in Table 3 (*P* = 0.168).

**Fig. 5.**
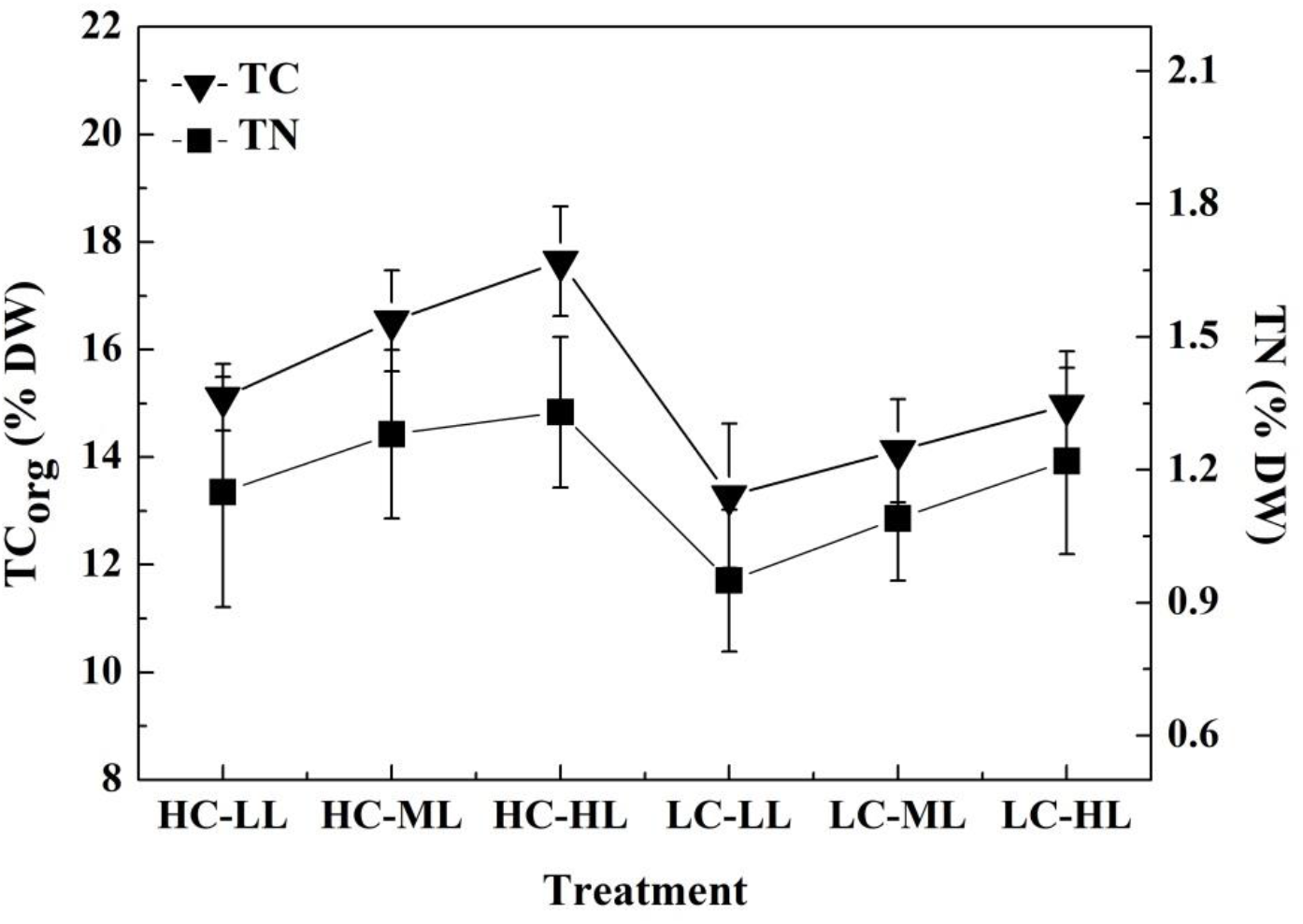
Variations in tissue total organic carbon (TC_org_, % DW) and nitrogen (TN, % DW) from the tissues of *Halimeda opuntia* (mean ± SD, n = 3) in six *p*CO_2_–light treatments.

### 3.5 Enzyme activities (external carbonic anhydrase and nitrate reductase)

The eCAA was significantly higher in CO_2_ enrichment than in ambient *p*CO_2_ conditions (Fig. 6; Table 3). Under 1400 ppmv *p*CO_2_, the eCAA of *H. opuntia* was 0.455–0.518 IU mg^−1^ FW, increasing by 13.35%–22.31% compared to that of the 400 ppmv *p*CO_2_. Increased light availability also displayed a positive role in regulating eCAA (*P* < 0.01) (Table 3). The NRA was affected by a significant interaction between elevated *p*CO_2_ and light intensities (Table 3). Similar to the eCAA, NRA increased in HC with values of 0.553–0.698 pg mg^−1^ FW, compared with values of 0.361–0.526 pg mg^−1^ FW in LC (Fig. 6). Moderately increasing light intensities positively enhanced the NRA by7.78%–26.22% and 30.75%–45.71% in HC and LC, respectively (Fig. 6).

**Fig. 6.**
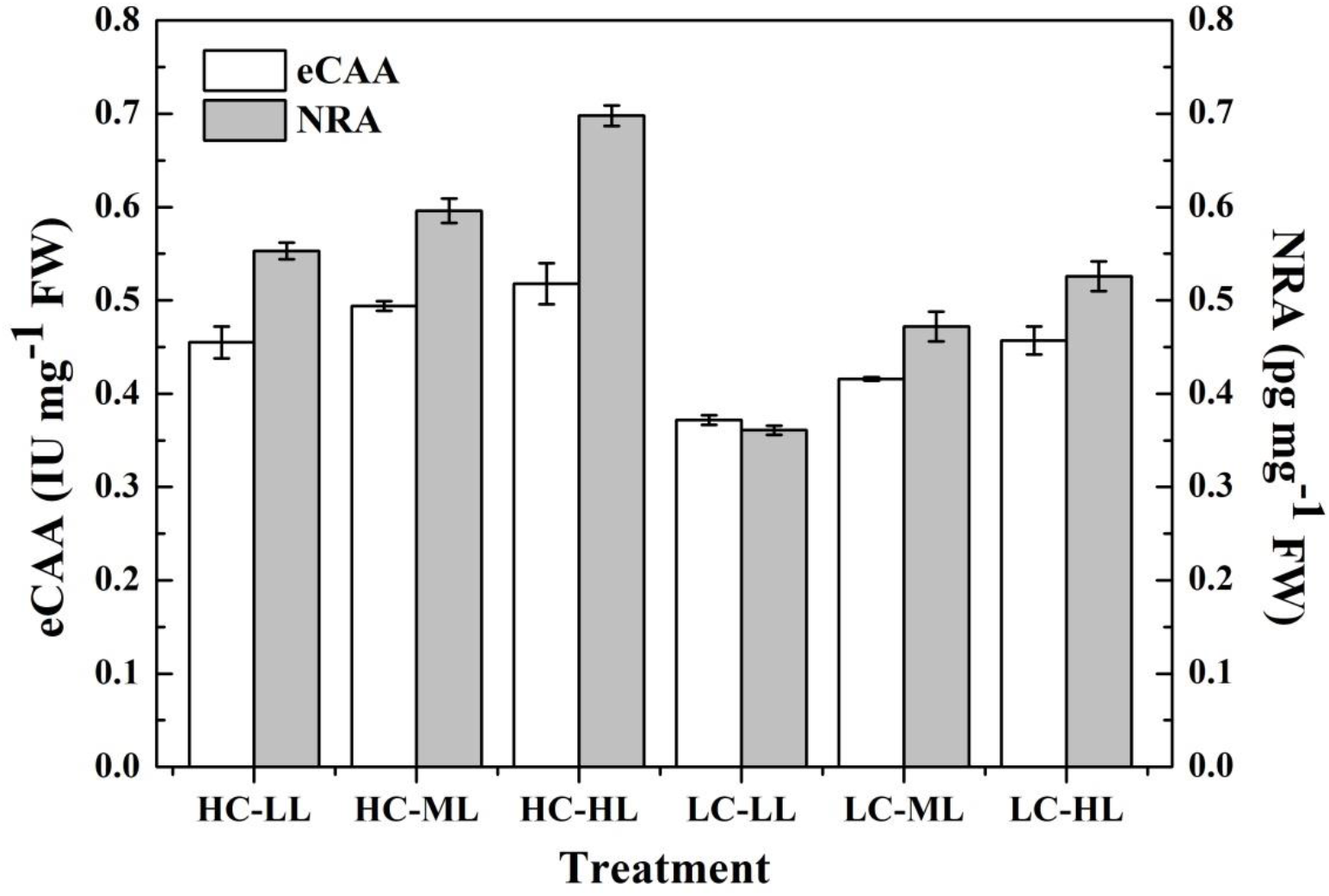
Mean (± SD, n = 3) external carbonic anhydrase activity (eCAA, IU mg^−1^ FW) and nitrate reductase activity (NRA, pg mg^−1^ FW) in six *p*CO_2_-light treatments.

### 3.6 Soluble carbohydrate, protein and free amino acid content

The SC profiles measured in *H. opuntia* at the end of the experiment were notably affected by elevated *p*CO_2_, light intensities and their interactions (*P* < 0.01) (Table 3). In HC, SC content ranged from 0.432 ± 0.021 μg mg^−1^ FW to 0.668 ± 0.014 μg mg^−1^ FW, with 12.21%–51.47% higher than in LC (0.385–0.441 μg mg^−1^ FW). Compared to LL conditions, ML and HL intensities (150–240 μmol photons m^−2^ s^−1^) SC content increased by 23.38%–54.63% and 12.73%–14.55% in HC and LC, respectively (Fig. 7a). Similar trends were documented for SP content in *H. opuntia*. The SP content in HC (0.347–0.547 μg mg^−1^ FW) was 11.58%–15.08% higher than in LC (0.311–0.478 μg mg^−1^ FW) (Fig. 7b). In addition, there was a positive effect on light availability on SP accumulation with an increase of 25.65%–57.64% and 21.86%–53.70% in HC and LC, respectively (Fig. 7b, Table 3). The ratios of SC/SP at HC levels were remarkably higher than in LC, and the lowest ratio of SC/SP value (0.923) occurred in LC-HL treatment (Fig. 7d). FAA content in *H. opuntia* was notably impacted by the conditions as shown in Table 3 (*P* < 0.01). Elevated *p*CO_2_ also significantly increased FAA content, which was 33.41%–54.74% higher than the thalli grown in ambient *p*CO_2_ (Fig. 7c).

**Fig. 7.**
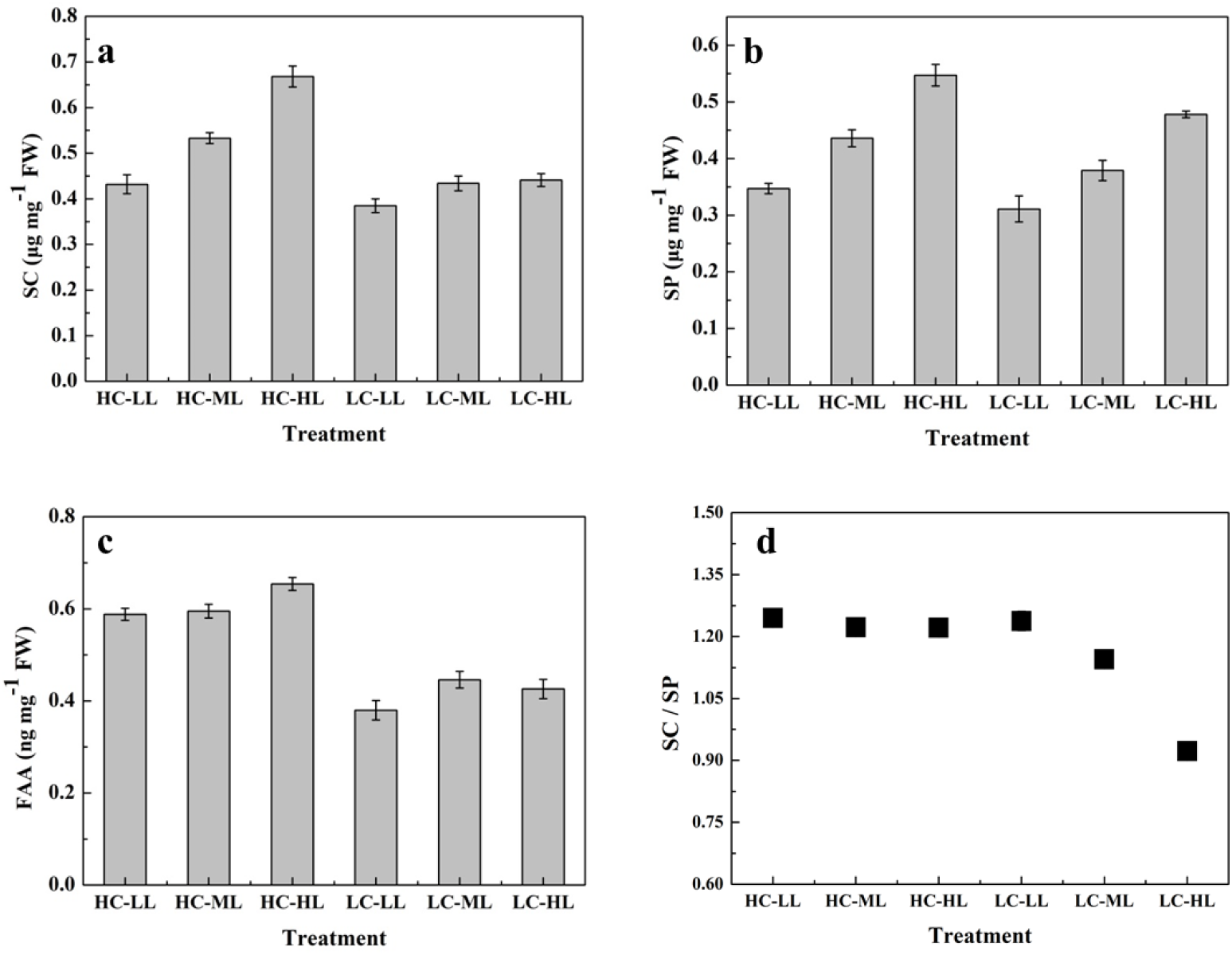
Variations in soluble carbohydrate (SC, μg mg^−1^ FW) (a), soluble protein (SP, μg mg^−1^ FW) (b), free amino acids (FAA, ng mg^−1^ FW) (c), and SC/SP (d) of *Halimeda opuntia* (mean ± SD, n = 3) in six *p*CO_2_–light treatments.

## 4. Discussion

The effects of CO_2_-induced OA have caused growing concerns regarding potential shifts in marine organisms, biological communities and coral reef ecosystems (Moodley et al., 2000; Feely et al., 2004; Zeebe et al., 2008). In common with previous studies, the data in the present study suggest that OA leads to deleterious impacts on growth rates, calcification and photosynthesis processes in calcifying macroalga species of *H. opuntia* (Fig. 1; Fig. 2). However, this study further revealed that photosynthetic products, such as SC, SP and FAA contents (Fig. 7), as well as relative enzymatic activities (Fig. 6), increased notably under higher light intensity, which could favor an enhanced performance in photosynthesis (Fig. 2-4), and tissue carbon and nitrogen accumulation (Fig. 5). The results indicate that moderately increased light availability could enhance the key anti-stress abilities of *H. opuntia* in response to OA. This conclusion is consistent with previous studies reported by Wei et al. (2019; 2020a), who suggested that OA and increased light influenced photosynthesis, calcification and other physiological performances of two *Halimeda* species (*H. cylindracea* and *H. lacunalis*) in opposing directions, and the positive influences of moderate increases in light availability could offset the adverse effects of the decline in pH caused by elevated *p*CO_2_.

In this study, *H. opuntia* performed lower *RGR* and calcification rate (*G*net) in ambient*p*CO_2_ conditions. Most of the previous work on *Halimeda* species (Teichberg et al., 2013; Hofmann et al., 2014; Campbell et al., 2016; Peach et al., 2017) and other marine calcifiers (Hofmann et al, 2008; Manzello, 2010; O’Donnell et al., 2010) under OA has focused on reduced growth, calcification and primary productivity. With CO_2_ enrichment, the components of dissolved inorganic carbon (DIC) drastically shifted and aragonite saturation state (Ω_Arag_) decreased (Table 2), which may contribute to the dissolution of calcium carbonate structures formed in algal tissues (Orr et al., 2005; Anderson et al., 2009). The decline in *G*_net_ presented here confirms the susceptibility of algal skeleton formation to slight changes in environmental pH values (Hofmann et al., 2014). Notably, enhancements of *RGR* and *G*_net_ were pronounced under increased light availability. As has been observed in other *Halimeda* species under OA, the calcification data provide evidence that increased light could be able to compensate for elevated *p*CO_2_ levels through the up-regulation of photosynthesis (Fig. 3) (Vásquez-Elizondo and Enríquez, 2016; Campbell et al., 2016). This is probably because algal calcification is energetically dependent on photosynthesis and could provide a high fraction of the energetic costs of the biomineralization process (Chisholm, 2000, 2003; Meyer et al., 2016). As calcification occurs in semi-enclosed inter-utricular spaces in *H. opuntia*, efficient photosynthesis aids crystal growth via CO_2_ removal and the elevation of intracellular pH (Campbell et al., 2016). Peach et al. (2017) also pointed out that increased photosynthesis may compensate for the adverse influences on algal calcification processes in response to OA, and contribute to skeletal CaCO_3_ precipitation.

Elevated *p*CO_2_ caused adverse impacts on thallus photosynthesis in *H. opuntia*, and the severity of this impact was observed under low light condition (30 μmol photons m^−2^ s^−1^), resulting in declines in *F*_v_/*F*_m_ and rETR (Fig. 2; Fig. 4a, b). A similar negative impact by elevated *p*CO_2_ on algal photosynthesis has been reported for other *Halimeda* species (Price et al., 2011; Sinutok et al., 2012; Compbell et al., 2015; Meyer et al., 2016). Price et al. (2011) found that *H. opuntia* had carbonate dissolution and a reduction in photosynthetic capacity after two weeks of exposure to *p*CO_2_ at 900 ± 90 ppmv. However, the specific reasons for these declines have not been concluded. According to the results of our 28-day experiments, it could be proposed that increasing dissolved CO_2_ in seawater reduced pH, which may depress the expression of CCMs in terms of DIC acquisition (Price et al., 2011). Although CCMs could enhance photosynthetic efficiency, they were energetic cost processes resulting from the activities of anhydrase to catalyze the hydration of CO_2_ and active ion pumping across membranes (Prins et al., 1982). As elevated *p*CO_2_ caused the degradation of CCM induction, algal photosynthesis majorly relied on passive CO_2_ diffusion, and thus the efficiency of photosynthetic carbon assimilation was critically limited. The maintenance of high proton-H^+^ permeability of the plasma membrane, which is essential for photosynthetic bicarbonate assimilation (Elzenga and Prins, 1989), declined at reduced external pH (Miedema and Prins, 1991). According to our results, the excess dissolved CO_2_ compromised both photophysiology and ΩArag for calcification in *H. opuntia*, indicating that OA may bring adverse synergistic depressions for this calcifying species. The metabolic suppression of photosynthesis and calcification activities suggests that either there is a decreased demand for ATP or that metabolism was actively suppressed as an acute survival response. This would potentially allow the reallocation of energy to a more immediate stress response, such as mucus production, which needs the maintenance of intracellular pH homeostasis and/or immune defense (Moya et al., 2012).

Concurrently, *H. opuntia* can resist the environmental stress caused by rapid seawater changes in OA by regulating pigment synthesis. A significant decline in Chl *a* content with elevated *p*CO_2_ (Fig. 4a) suggested pigment degradation or/and increases in light stress (Sinutok et al., 2012). However, carotenoid content, especially in low light conditions were remarkably accumulated and may function to take full use of light energy during periods of lower irradiance (Boardman, 1977; Teichberg et al., 2013). Based on the photosynthetic performance, although values of *F*_v_/*F*_m_ (0.665–0.704) declined in elevated *p*CO_2_, it remained at a relatively normal level after 28-day incubation, which indicates that the effects of a moderate increase in light availability could offset the severe impacts of low pH (Wei et al., 2020a). The rapid light curves for rETR were significantly enhanced by increasing light intensities as displayed in Fig. 2. Vásquez-Elizondo and Enríquez (2016, 2017) suggested that changes in the internal carbon and nitrogen partitioning of the algal thallus induced by elevated *p*CO_2_ (as shown in Fig. 5 and Fig. 7 in this study) affected the optical properties of the antenna size of the photosystems. This new organization of the photosynthetic machinery indicates that *H. opuntia* accommodated light variations and elevated *p*CO_2_ conditions (Wei et al., 2020a).

Another striking anti-stress ability exhibited by *H. opuntia* in response to OA was observed in TCorg and TN accumulation in algal tissues (Fig. 5). Similar conclusions have been reached in some other comparative studies and it was proposed that these C and N accumulations resulted from characterizations of enzymatic-driven activities, especially for the eCAA and NRA (Hofmann et al., 2014; Chen et al., 2017; 2018). In the present study, CO_2_ enrichment as well as increased light intensities positively enhanced eCAA of *H. opuntia*, which catalyzes CO_2_ and HCO_3_^−^ interconversion and ensures that the substrate for Rubisco remains at a normal equilibrium at the site of carboxylation under elevated *p*CO_2_ (Moya et al., 2012; Vidal-Dupiol et al., 2013; Hofmann et al., 2014). As a result of increasing dissolved CO_2_ in seawater (Table 2), concentration gradients between the seawater environment and algal cytoplasm may function to favor the diffusion of CO_2_ and HCO_3_^−^ (Moroney et al., 1985; Zou and Gao, 2004). Thus, the SC synthesis in *H. opuntia* was enhanced by the positive effects of CA activity and elevated *p*CO_2_ (Fig. 7a), which resulted in an improvement in the irradiance harvesting capacity to absorb light and the up-regulation of the photosynthetic electron transport chain (Van Oijen et al., 2004). Moreover, CO_2_ enrichment in this study was considered to significantly increase both NRA and TN accumulation in *H. opuntia* (Fig. 5; Fig. 6), which is consistent with the conclusions reported by Chen et al. (2016; 2017) and Wei et al. (2020a). Indeed, a positive correlation between increased light availability and higher TN content was observed during the experimental period, even though it was not statistically significant (*P* = 0.168) (Table 4). These results indicate that higher light might cause metabolic modifications in *H. opuntia* to mitigate OA stress, which is confirmed by both the higher photosynthetic rates and soluble organic osmolytes (Fig.7).

It has been suggested that soluble organic molecules may play roles in buffering metabolic depression to cope with OA challenges (Xiong et al., 2002; Sun et al., 2013; Chen et al., 2017; 2018; Wei et al., 2020a). Light-mediated changes in cellular biochemical composition were associated with the acclimation of algal growth and photosynthesis (Zou and Gao, 2013). Under elevated *p*CO_2_, SC and SP contents were remarkably higher than those of *H. opuntia* tissues under control *p*CO_2_ as displayed in Fig. 7. As the SC content increased, the cytoplasmic concentration was promoted whereas the permeability of the plasma membrane declined (Xiong et al., 2002). These modifications contribute to membrane integrity and ensure cellular activities function well, and could provide the physiological basis for the resistance of adverse external environments by algal cells (Sun et al., 2013; Chen et al., 2017; Wei et al., 2020a). Such shifts in metabolic performance may result from the enhancement of internal and external CA activities that facilitates the conversion of CO_2_ and HCO_3_^−^ to satisfy photosynthetic carbon demand (De Beer and Larkum, 2001). Thus, increased SC contents in *H. opuntia* favored the enhancement of resilience against OA stress. The SP content was another important indicator of plant metabolism (Sun et al., 2015). Based on the results (Fig. 7b), an obvious trend in CO_2_-induced promotion of SP content was observed in CO_2_ enrichment treatments. Yang et al. (2009) pointed out that a higher SP accumulation works in increasing key enzyme activities for plant photosynthesis and accelerating substance and energy metabolism. Furthermore, consistent with previous studies (Rizhsky et al., 2004; Fougère et al., 2008; Wei et al., 2020a), it is reasonable to suggest that the stimulation of FAAs might increase the anti-oxidative capacity of *H. opuntia*. Wei et al. (2020a) demonstrated that under elevated *p*CO_2_ (1600 ppmv), proline contents, one of the FAAs in both *H. cylindracea* and *H. lacunalis,* were two- to four-fold higher than those in ambient *p*CO_2_ (400 ppmv). These results suggested that FAA contents were largely secreted and work in protecting cellular structures, scavenging reactive oxygen species (ROS) and related enzyme systems (Verma, 1999; Xiong et al., 2002).

Notably, higher light intensities and eCAA could promote photosynthesis processes, which in turn increased photosynthetic products and SC accumulation to improve flexibility and plasticity in response to OA. As SP and FAA synthesis largely depended on photosynthesis and SC content the assimilation of N in macroalgae required both carbon skeletons and energy (Chen et al., 2016; 2017). Moreover, in the present study, the up-regulation of eCAA and NRA was significantly promoted by CO_2_ levels and/or increased light availability, which favored inorganic carbon and nitrogen uptake and assimilation (Losada and Guerrero, 1979; Syrett, 1981). Taken together, these appropriate physiological changes of soluble organic molecules (SC, SP and FAA) describe an anti-stress ability in which relatively high light intensity increases in metabolic functioning combined with enzymatic activities can mitigate OA stress in *Halimeda* species.

## 5. Conclusion

Deteriorating marine conditions caused by human activities threatens the diversity, distribution and biomass of coral reef ecosystems, including calcifying macroalgae. According to the results of the current study, the broader evidence indicates that OA exhibits an adverse role in influencing growth, calcification, photosynthesis and other physiological performance of *H. opuntia*. However, our results also indicate that increased light availability can enhance anti-stress abilities by the accumulation of soluble organic molecules, especially for SC, SP and FAA, in combination with metabolic enzyme-driven activities to alleviate OA stress. The results offer a more complete understanding of whether marine calcifiers have the physiological plasticity to compensate for the effects of OA and whether they will continue to build skeletons under future CO_2_ conditions.

## Acknowledgements

This research was supported by the Special Research Assistant Grant program of the Chinese Academy of Sciences, the National Natural Science Foundation of China (42006129), Key Special Project for Introduced Talents Team of Southern Marine Science and Engineering Guangdong Laboratory (Guangzhou) (GML2019ZD0404) and the Strategic Priority Research Program of the Chinese Academy Sciences (XDA13020203). Thanks are also due to editors and anonymous reviewers for their valuable comments and suggestions.

